# Odorant receptors from *Culex quinquefasciatus* and *Aedes aegypti* sensitive to floral compounds

**DOI:** 10.1101/721878

**Authors:** Fangfang Zeng, Pingxi Xu, Walter S. Leal

## Abstract

Mosquitoes rely heavily on the olfactory system to find a host for a bloodmeal, plants for a source of energy and suitable sites for oviposition. Here, we examined a cluster of 8 odorant receptors (ORs), which includes one OR, CquiOR1, previously identified to be sensitive to plant-derived compounds. We cloned 5 ORs from *Culex quinquefasciatus* and 2 ORs from *Aedes aegypti*, ie, CquiOR2, CquiOR4, CquiOR5, CquiOR84, CquiOR85, AaegOR14, and AaegOR15 and then deorphanized these receptors using the *Xenopus* oocyte recording system and a large panel of odorants. 2-Phenylethanol, phenethyl formate, and phenethyl propionate were the best ligands for CquiOR4 somewhat resembling the profile of AaegOR15, which gave the strongest responses to phenethyl propionate, phenethyl formate, and acetophenone. In contrast, the best ligands for CquiOR5 were linalool, PMD, and linalool oxide. CquiOR4 was predominantly expressed in antennae of nonblood fed female mosquitoes, with transcript levels significantly reduced after a blood meal. 2-Phenylethanol showed repellency activity comparable to that of DEET at 1%. RNAi experiments suggest that at least in part 2-phenylethanol-elicited repellency is mediated by CquiOR4 activation.

## 1. Introduction

Mosquitoes rely on the olfactory system to find plants as a source of carbohydrates, hosts for bloodmeals, and oviposition sites. Because pathogens might be transmitted during a bite by an infected mosquito, there is understandably a great deal of interest in unraveling the olfactory aspects of human-mosquito interactions (Bradshaw et al., 2018) to explore ways of reducing mosquito bites. However, plant nectar sources are often essential for mosquitoes because they increase mosquito life span and reproductive capacity (Lahondère et al., 2019) and long-living mosquitoes are more dangerous (Tan et al., 2019). Therefore, understanding how mosquitoes find plants/flowers is also important for reducing the transmission of vector-borne diseases.

Previously, we have identified generic and plant kairomone sensitive odorant receptors (ORs) from the Southern house mosquito, *Culex quinquefasciatus* (Xu et al., 2013). One of these ORs, CquiOR1, belongs to a cluster of 6 ORs from the Southern house mosquito and 2 ORs from the yellow fever mosquito, *Aedes (=Stegomyia) aegypti* (Fig. S1). Specifically, CquiOR2, CquiOR4, AaegOR14, AaegOR15, CquiOR5, CquiOR84, and CquiOR85. Of note, CquiOR2 (VectorBase, CPIJ0000542) is not the previously reported oviposition attractant-detecting OR2 (Pelletier et al., 2010), which has been renamed CquiOR121 (CPIJ014392) (Leal et al., 2013). We cloned CquiOR2 (Genbank, MG280964) and the other ORs in this cluster (CquiOR4, MG280965, AaegOR14, MN227017, AaegOR15, MN227018, CquiOR5, MG280966, CquiOR84, MN227015, and CquiOR85, MN227016) and deorphanized them. Here, we report that these receptors, particularly CquiOR4, CquiOR5, and AaegOR15 are very sensitive to plant-derived compounds, including repellents. CquiOR4, for example, which is very specific to female antennae, with high and low transcript levels in nonblood fed and blood-fed mosquitoes, respectively, showed a robust response to the natural repellent 2-phenylethanol. Repellency activity elicited by 2-phenylethanol reduced significantly in CquiOR4-dsRNA-treated mosquitoes, but it was unchanged when these mosquitoes were tested against DEET, which is detected with another receptor (Xu et al., 2014).

## 2. Materials and Methods

### 2.1. Insect Preparations and Sample Collection

Mosquitoes used in this study were from a laboratory colony of *Cx. quinquefasciatus* originating from adult mosquitoes collected in Merced, CA in the 1950s and kept at the Kearney Agricultural Research Center, University of California, Parlier, CA. Specifically, we used mosquitoes from the Davis colony, which was initiated about 8 years ago with mosquitoes from the Kearney colony. In Davis, mosquitoes were maintained at 27±1°C, 75±5% relative humidity, and under a photoperiod of 12:12 h.

### 2.2. OR Cloning

Total RNA samples were extracted from one thousand 4-7 day-old *Culex* female antennae with TRIzol reagent (Invitrogen, Carlsbad, CA). Antennal cDNA was synthesized from 1 µg of antennal total RNA from each species using iScript cDNA synthesis kit (Bio-Rad) according to the manufacturer’s instructions (Clontech, Mountain View, CA). Total RNA was extracted from *Aedes* mosquitoes provided by Dr. Anthon J. Cornel. To obtain full-length coding sequences of CquiOR2, CquiOR4, CquiOR5, CquiOR84, CquiOR85, AaegOR14, and AaegOR15, PCRs were performed using the following gene-specific primers containing restriction endonuclease sites (XmaI and XbaI) and Kozak motif (acc):

InFu-CqOR2-F: AGATCAATTC<u>CCCGGG</u>accATGAGGTTCGCCCCGCTC

InFu-CqOR2-R: TCAAGCTTGC<u>TCTAGA</u>TCAAATGCTATCCTTTAAAATCACA

InFu-CqOR4-F: AGATCAATTC<u>CCCGGG</u>accATGAAATCCCACAGTCCCCTCAA

InFu-CqOR4-R: TCAAGCTTGC<u>TCTAGA</u> TTACAACCTCTCCTTCAGCACGACA

InFu-CqOR5-F: AGATCAATTC<u>CCCGGG</u>acc ATGAAATTCTACGAGCTCCGCG

InFu-CqOR5-R: TCAAGCTTGC<u>TCTAGA</u> TTATGAATGCATCAATCGCTCCCT

InFu-CqOR84-F: GATCAATTC<u>CCCGGG</u>accATGGAGTTCCTGGCCGC

InFu-CqOR84-R: CAAGCTTGC<u>TCTAGAT</u>TAGTTCACTCCCTGCAGTCG

InFu-CqOR85-F: AGATCAATTC<u>CCCGGG</u>accATGGAGTTCCTGGCCGCC

InFu-CqOR85-R: TCAAGCTTGC<u>TCTAGA</u>TTAGATCACTCCCTGCAGTCGTTC

InFu-AaOR14-F: GATCAATTC<u>CCCGGG</u>accATGAACTACTTTGAGCTA

InFu-AaOR14-R: CAAGCTTGC<u>TCTAGA</u>TTACAAGTAATCCTTAAGCACC

InFu-AaOR15-F: GATCAATTC<u>CCCGGG</u>accATGAAGTACTTTGAGCT

InFu-AaOR15-R: CAAGCTTGC<u>TCTAGA</u>TTACAACTGATCCTTTAGTACAACGT

PCR products were purified by a QIAquick gel extraction kit (Qiagen) and then subcloned into pGEMHE vector with In-Fusion HD cloning Kit (Clontech, Mountain View, CA). After transformation, plasmids were extracted using the QIAprep Spin Miniprep kit (Qiagen) and sequenced by ABI 3730 automated DNA sequencer at Davis Sequencing (Davis, CA) for confirmation.

### 2.3. Electrophysiology

Two-electrode voltage-clamp technique (TEVC) was performed as previously described (Xu et al., 2014). Briefly, the capped cRNAs were synthesized using pGEMHE vectors and mMESSAGE mMACHINE T7 Kit (Ambion). Purified OR cRNAs were resuspended in nuclease-free water at 200 ng/mL and microinjected with the same amount of CquiOrco (or AaegOrco) cRNA into *Xenopus laevis* oocytes in stage V or VI (purchased from EcoCyte Bioscience, Austin, TX). Then, the oocytes were kept at 18°C for 3-7 days in modified Barth’s solution [NaCl 88 mM, KCl 1 mM, NaHCO_3_ 2.4 mM, MgSO_4_ 0.82 mM, Ca(NO_3_)2 0.33 mM, CaCl_2_ 0.41 mM, HEPES 10 mM, pH 7.4] supplemented with 10 mg/mL of gentamycin, 10 mg/mL of streptomycin. Odorant-induced currents at holding potential of −80 mV were collected and amplified with an OC-725C amplifier (Warner Instruments, Hamden, CT), low-pass filtered at 50 Hz and digitized at 1 kHz. Data acquisition and analysis were carried out with Digidata 1440A and pCLAMP 10 software (Molecular Devices, LLC, Sunnyvale, CA). The panel of odorants included the following compounds: 1-butanol, 1-pentanol, 1-hexanol, 1-heptanol, 1-octanol, 1-nonanol, 1-dodecanol, 2,3-butanediol, 2-butoxyethanol, 3-methyl-1-butanol, 2-hexen-1-ol, 3-hexen-1-ol, 1-hexen-3-ol, 1-hepten-3-ol, 3-octanol, 1-octen-3-ol, 2-octanol, 2-butanol, 2-nonen-1-ol, 2-pentanol, 4-methylcyclohexanol, 1-hexadecanol, 3-pentanol, 3-methyl-2-butanol, 3-methyl-2-buten-1-ol, 2-methyl-3-buten-2-ol, propanal, butanal, pentanal, isovaleraldehyde, hexanal, (E)-2-methyl-2-butenal, heptanal, octanal, nonanal, decanal, undecanal, 1-dodecanal, (E)-2-hexenal, (Z)-8-undecenal, (E)-2-heptenal, (E)-2-nonenal, phenylacetaldehyde, 2,4-hexadienal, furfural, benzaldehyde, α-hexylcinnamaldehyde, methyl acetate, ethyl acetate, propyl acetate, butyl acetate, pentyl acetate, hexyl acetate, heptyl acetate, octyl acetate, nonyl acetate, decyl acetate, methyl propionate, ethyl propionate, methyl butyrate, ethyl butanoate, methyl hexanoate, ethyl hexanoate, ethyl 3-hydroxyhexanoate, ethyl 3-hydroxybutanoate, ethyl linoleate, phenyl propanoate, phenethyl propionate, ethyl 2-(E)-4-(Z)-decadienoate, (E)-2-hexenyl acetate, (Z)-3-hexenyl acetate, (E)-2-hexenyl acetate, ethyl lactate, phenyl isobutyrate, eugenyl acetate, methyl salicylate, ethyl stearate, methyl myristate, isopropyl myristate, palmitic acid methyl ester, 1-octen-3-yl acetate, isopentyl acetate, ethyl phenylacetate, geranyl acetate, octadecyl acetate, acetylacetone, 2-butanone, 2-heptanone, geranylacetone, 6-methyl-5-hepten-2-one, 5-methyl-2-hexanone, 2,3-butanedione, 3-hydroxy-2-butanone, 2-pentanone, 2-hexanone, 2-octanone, 2-undecanone, 2-tridecanone, 2-nonanone, 1-octen-3-one, cyclohexanone, acetophenone, γ-valerolactone, γ-hexalactone, γ-octalactone, γ-decalactone, γ-dodecalactone, p-coumaric acid, isovaleric acid, dodecanoic acid, (±)-lactic acid, ethanoic acid, propanoic acid, butanoic acid, isobutyric acid, 2-oxobutyric acid, pentanoic acid, 2-oxovaleric acid, myristic acid, palmitoleic acid, oleic acid, hexanoic acid, (E)-2-hexenoic acid, 5-hexanoic acid, (E)-3-hexenoic acid, heptanoic acid, octanoic acid, nonanoic acid, decanoic acid, n-tridecanoic acid, linoleic acid, ammonia, trimethylamine, propylamine, butylamine, pentylamine, hexylamine, heptylamine, octylamine, 1,4-diaminobutane, cadaverine, 1,5-diaminopentane, phenol, 2-methylphenol, 3-methylphenol, 4-methylphenol, 4-ethylphenol, 3,5-dimethylphenol, 2,3-dimethylphenol, 2,4-dimethylphenol, 2,5-dimethylphenol, 2,6-dimethylphenol, 3,4-dimethylphenol, guaiacol, 2-methoxy-4-propylphenol, 2-phenoxyethanol, 1,2-dimethoxybenzene, benzyl alcohol, 2-phenylethanol, 1-phenylethanol, phenylether, (S)-(-)-perillaldehyde, fenchone, thujone, camphor, α-terpinene, γ-terpinene, (—)-menthone, menthyl acetate, limonene, linalyl acetate, α-humulene, linalool oxide, geraniol, nerol, thymol, (±)-linalool, eucalyptol, citral, eugenol, α-pinene, ocimene, (±)-citronellal, α-phellandrene, nerolidol, jasmine, menthol, carvone, cymene, terpinolene, ß-myrcene, (+)-δ-cadinene, (+)-limonene oxide, (E,E)-farnesol, (E,E)-farnesyl acetate, farnesene, α-methylcinnamaldehyde, cinnamyl alcohol, α-terpineol, citronellol, (E)-cinnamaldehyde, (-)-caryophyllene oxide, ß-caryophyphyllene, carvacrol, terpinen-4-ol, 7-hydroxycitronellal, pyridine, pyrrolidine, 2-pyrrolidinone, indole, 3-methylindole, isoprene, 5-isobutyl-2,3-dimethylpyrazine, dibutyl phthalate, dimethyl phthalate, phenethyl formate, benzyl formate, 2-acetylthiophene, methyl disulfide, 2-ethyltoluol, 2-methyl-2-thiazoline, methyl anthranilate, 4,5-dimethylthiazole, p-mentane-3,8-diol (PMD), N-(2-isopropyl-phenyl)-3-methyl-benzamide, and N,N-diethyl-3-methylbenzamide (DEET).

### 2.4. dsRNA Synthesis

Double-strand RNA (dsRNA) of CquiOR4, CquiOR5 and β-galactosidase were synthesized by in vitro transcription from PCR product using the MEGAscript T7 transcription kit (Ambion, Austin, TX). PCR was performed using plasmids containing the target genes as DNA template with the following gene-specific primers that included T7 promoter sequence (underlined);

Cquiβ-gal-F: 5′-<u>TAATACGACTCACTATAGGG</u>AATGGTTCAGGTCGAAAACG-3′ and Cquiβ-gal-R: 5′-<u>TAATACGACTCACTATAGGG</u>CCGCCTCGTACAAAACAAGT-3′.

CquiOR4-F: 5’ <u>AATACGACTCACTATAGGG</u>GTGCGTGAAACTGTTCGGA-3’

CquiOR4-R: 5’ <u>TAATACGACTCACTATAGGG</u>GCGAGTGTCCAGCCCGTA-3’

CquiOR5-1-F: 5’<u>TAATACGACTCACTATAGGG</u>ATGAAATTCTACGAGCTC

CquiOR5-1-R: 5’<u>TAATACGACTCACTATAGGG</u>AGTAATAAGCATGACATG

CquiOR5-2-F: 5’<u>TAATACGACTCACTATAGGG</u>CCGATGCACTTGCTGTTAG

CquiOR5-2-R: 5’<u>TAATACGACTCACTATAGGG</u>TTATGAATGCATCAATCGCT

Large scale dsRNAs were purified with MEGAclear kit (Ambion, Austin, TX) and precipitated with 5 M ammonium acetate to yield 4-5 μg/μl of CquiOR4&5-dsRNA.

### 2.5. dsRNA Microinjection

Female pupae (0-d-old) were collected in plastic cups filled with distilled water and kept on ice for 15 min. The sharp end of a yellow pipette tip was cut diagonally to make a stage to hold a pupa. Forty-five nanograms of dsRNAs in 9.2 nL volume were injected in the dorsal membrane close to the base of the trumpet using a NanoLiter 2000 inject (World Precision Instruments). The injected pupae were put in new plastic cups with distilled water and kept at 27 °C. Newly emerged adults were supplied with sugar water (10% wt/vol), and newly emerged males (ratio 1:1) were released into the cage for mating.

### 2.6. Quantitative Analysis of Transcription Levels

For tissue expression pattern analysis, each type of tissue (antennae, maxillary palps, proboscis, and legs) from three hundred non blood-fed female mosquitoes (4-5 days old) and antennae from three hundred blood-fed female mosquitoes were dissected and collected in TRIzol reagent (Invitrogen, Carlsbad, CA) on ice using a stereomicroscope (Zeiss, Stemi DR 1663, Germany). For dsRNA treated mosquitoes, thirty pairs of antennae were dissected from each group and collected in 50% (vol/vol) ethanol diluted in DEPC-treated water on ice using a stereomicroscope. Total RNAs were extracted, and cDNAs were synthesized using iScript Reverse Transcription Supermix for RT-qPCR according to the manufacturer’s instructions (Bio-Rad). Real-time quantitative PCR (qPCR) was carried out by using a CFX96 Touch^TM^ Real-Time PCR Detection System (Bio-Rad) and SsoAdvanced SYBR Green Supermix (Bio-Rad). CquiRPS7 gene was used as a reference.

The following pairs of detection primers were designed with Primer 3 program (frodo.wi.mit.edu/):

CquiRPS7-Fw: 5’- ATCCTGGAGCTGGAGATGA −3’;

CquiRPS7-Rv: 5’- GATGACGATGGCCTTCTTGT −3’;

CquiOR4-qPCR-Fw: 5’- CTTTACCTCGGAGACCACCA-3’

CquiOR4-qPCR-Rv: 5’-ACCGGTAGCTTGATCCAGTG-3’

CquiOR5-qPCR-Fw: 5’-TTACTCCGATGCACTTGCTG-3’

CquiOR5-qPCR-Rv: 5’-TCTCGCAAAGTTGATCCAGA-3’

qPCR was performed with three biological replicates, and each of them was replicated three times (three technical replicates per biological replicate); data were analyzed using the 2^−ΔΔCT^ method.

### 2.7. Surface-Landing and Feeding Assay

The bioassay arena was modified from our surface-landing assay (Syed and Leal, 2008) initially designed to mimic a human arm without odors or humidity. CO_2_ at 50 mL/min was added to activate female mosquitoes, and blood was provided as both an attractant and a reward. In short, two 50-mL Dudley bubbling tubes, painted internally with a black hobby and craft enamel (Krylon, SCB-028), were held in a wooden board (30 × 30 cm), 17 cm apart from each end and 15 cm from the bottom. The board was attached to the frame of an aluminum collapsible field cage (30.5 × 30.5 × 30.5 cm; Bioquip). Two small openings were made 1 cm above each Dudley tube to hold two syringe needles (Sigma-Aldrich, 16-gauge, Z108782) to deliver CO_2_. To minimize the handling of mosquitoes, test females were kept inside collapsible field cages since the latest pupal stage. These female cages had their cover modified for behavioral studies. A red cardstock (The Country Porch, GX-CF-1) was placed internally at one face of the cage, and openings were made in the cardboard and cage cover so the cage could be attached to the wooden board with the two Dudley tubes and CO_2_ needles projecting inside the mosquito cage 6 and 3cm, respectively. Additionally, windows were made on the top and the opposite end of the red cardstock for manipulations during the assays and a video camera connection, respectively. The mosquito cage housing 30–50 test females was connected to the platform holding the Dudley tubes at least 2 h before bioassays. At least 10 min before the assays, water at 38°C started to be circulated with a Lauda’s Ecoline water bath, and CO^2^ at 50 mL/min was delivered from a gas tank just at the time of the behavioral observations. Sample rings were prepared from strips of filter papers 25 cm long and 4 cm wide and hung on the cardstock wall by insect pins to make a circle around the Dudley tubes. Cotton rolls (iDental, 1 × 3 cm) were loaded with 100 μL of defibrinated sheep blood purchased from the University of California, Davis, VetMed shop and placed between a Dudley tube and a CO_2_ needle. For each run one paper ring was loaded with 200 μL of hexane (control) and the other with 200 μL tested compounds at a certain concentration in hexane. The solvent was evaporated for 1–2 min, blood-impregnated cotton plugs, and filter paper rings were placed in the arena, CO_2_ was started, and the assays were recorded with a camcorder equipped with Super NightShot Plus infrared system (Sony Digital Handycan, DCR-DVD 810).

During the assay, the arena was inspected with a flashlight whose lens was covered with a red filter. After 5 min, the number of females that landed and continued to feed on each side of the arena was recorded. Insects were gently removed from the cotton rolls, and the assays were reinitiated after rotation of sample and control. Thus, repellency for each set of test mosquitoes was measured with the filter paper impregnated with the same sample at least once on the left and once on the right side of the arena. After three runs, filter paper strips and cotton plugs were disposed of, and new loads were prepared.

### 2.8. Phylogenetic Analysis of Mosquito ORs

Amino acid sequences of ORs from *Cx. quinquefasciatus, Ae. aegypti*, and *An. gambiae* were aligned by Clustal Omega (https://www.ebi.ac.uk/Tools/msa/clustalo/) with the generated FASTA file being used as an entry for phylogenetic analysis in MEGA7 (Kumar et al., 2016). Parameters were set as following: bootstrap method: 1000; p-distance; pairwise deletion. The generated tree was imported into iTOL (https://itol.embl.de/).

### 2.9. Graphic Preparations and Statistical Analysis

Graphic illustrations were prepared with Prism 8 (GraphPad, La Jolla, CA). The number of mosquitoes in the treatment (T) and control (C) side of the arena were transformed into % protection, P% = (1-[T/C]) × 100, according to WHO (WHO, 2009) and EPA (EPA, 2010) recommendations. Data that passed the Shapiro-Wilk normality test were analyzed with the two-tailed, unpaired *t*-test; otherwise, data were analyzed with the Mann-Whitney test. All data are expressed as mean ± SEM.

## 3. Results and Discussion

### 3.1. Deorphanization of Clustered *Culex* and *Aedes* ORs

We cloned the cDNAs for CquiOR2, CquiOR4, CquiOR5, CquiOR84, CquiOR85, AaegOR14, and AaegOR15, which are clustered (Fig. S1) along CquiOR1 (Xu et al., 2013). Then, each OR was separately co-expressed along with its coreceptor in *Xenopus* oocytes for deorphanization. We expected that the odorant profiles for these receptors would resemble that of CquiOR1 (Xu et al., 2013), but found marked differences. For example, three floral compounds elicited robust currents on CquiOR4/CquiOrco-expressing oocytes. Specifically, these oocytes were very sensitive to 2-phenylethanol, phenethyl formate, and, propionate (Fig. 1). To obtain more information about the sensitivity of the CquiOR4 + CquiOrco receptor, we performed concentration-response analyses for the three best ligands. Currents elicited by 2-phenylethanol were already saturated at the normal screening dose of 1 mM. Thus, we performed these analyses with concentrations in the range of 0.1 µM to 0.1 mM (Fig. 2). 2-Phenylethanol was indeed the most potent of the compounds in our panel, activating CquiOR4+CquiOrco with EC50 of 28 nM.

**Fig. 1.**
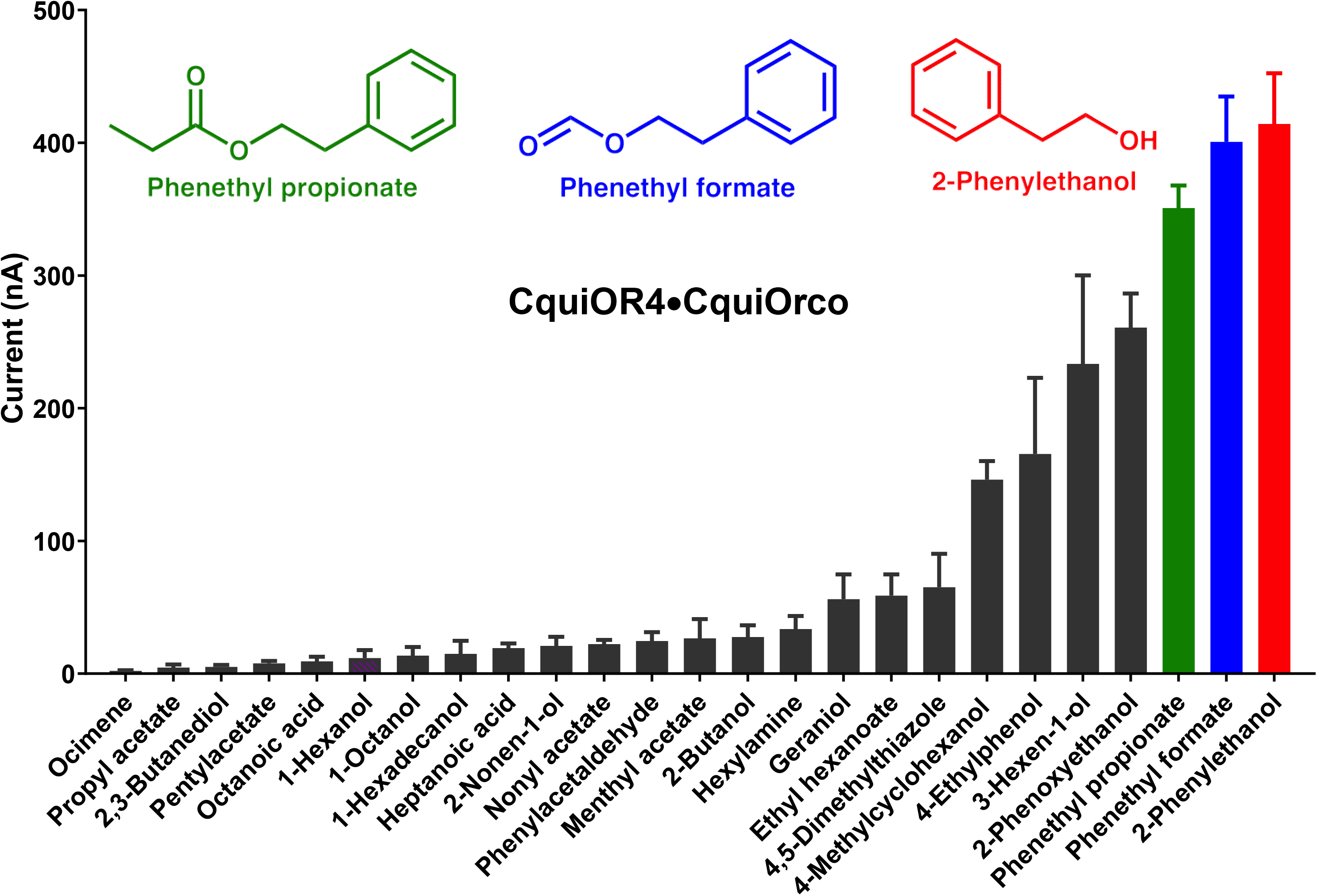
Quantification of current responses from oocytes expressing CquiOR4 and CquiOrco. All compounds in our panel were tested at 1 mM dose (n = 3). For clarity, bars are displayed in increasing order of response (Mean ± SEM) and by omitting compounds that did not elicit a response. The chemical structures of the three major ligands are displayed on the top of the graphic in colors corresponding to their bars.

**Fig. 2.**
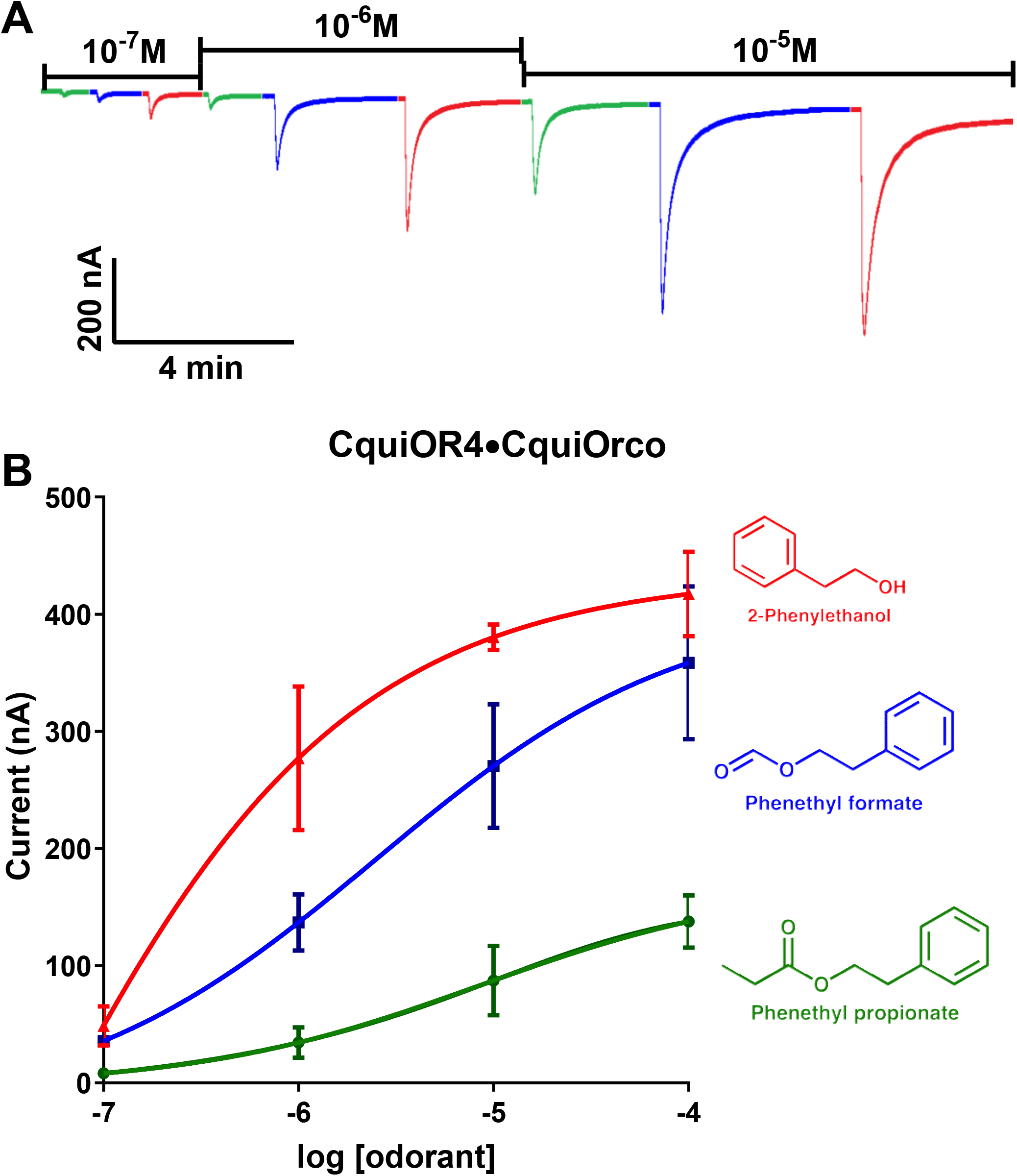
Concentration-dependent responses of CquiOR4⋅CquiOrco-expressing oocytes to three major odorants. (A) Traces obtained with a single oocyte challenged with 2-phenylethanol (red), phenethyl formate (strawberry) and phenethyl propionate (green) at the doses of 10^−7^ to 10^−5^ M. (B) Dose-dependent responses obtained with three oocytes and three replicates for each ligand. Responses (Mean ± SEM) were not normalized.

CquiOR5 + CquiOrco receptor showed a quite different profile, with the three best ligands being linalool, p-methane-3,8-diol (PMD) and linalool oxide (Fig. 3). Although the responses were not as robust as those elicited by the best ligands for CquiOR4 + CquiOrco, they were dose-dependent (Fig. S2). Linalool activated CquiOR5/CquiOrco-expressing oocytes with an EC50 of 574 nM. By contrast, no compounds in our panel elicited relevant currents in CquiOR2/CquiOrco-expressing oocytes, except for isopentyl acetate (Fig. S3A). On the other hand, CquiOR84/CquiOrco-expressing oocytes responded with large currents when challenged with N-(2-isopropyl-phenyl)-3-methyl-benzamide or PMD (Fig. S3B). Similar responses were recorded with CquiOR85 + CquiOrco receptor.

**Fig. 3.**
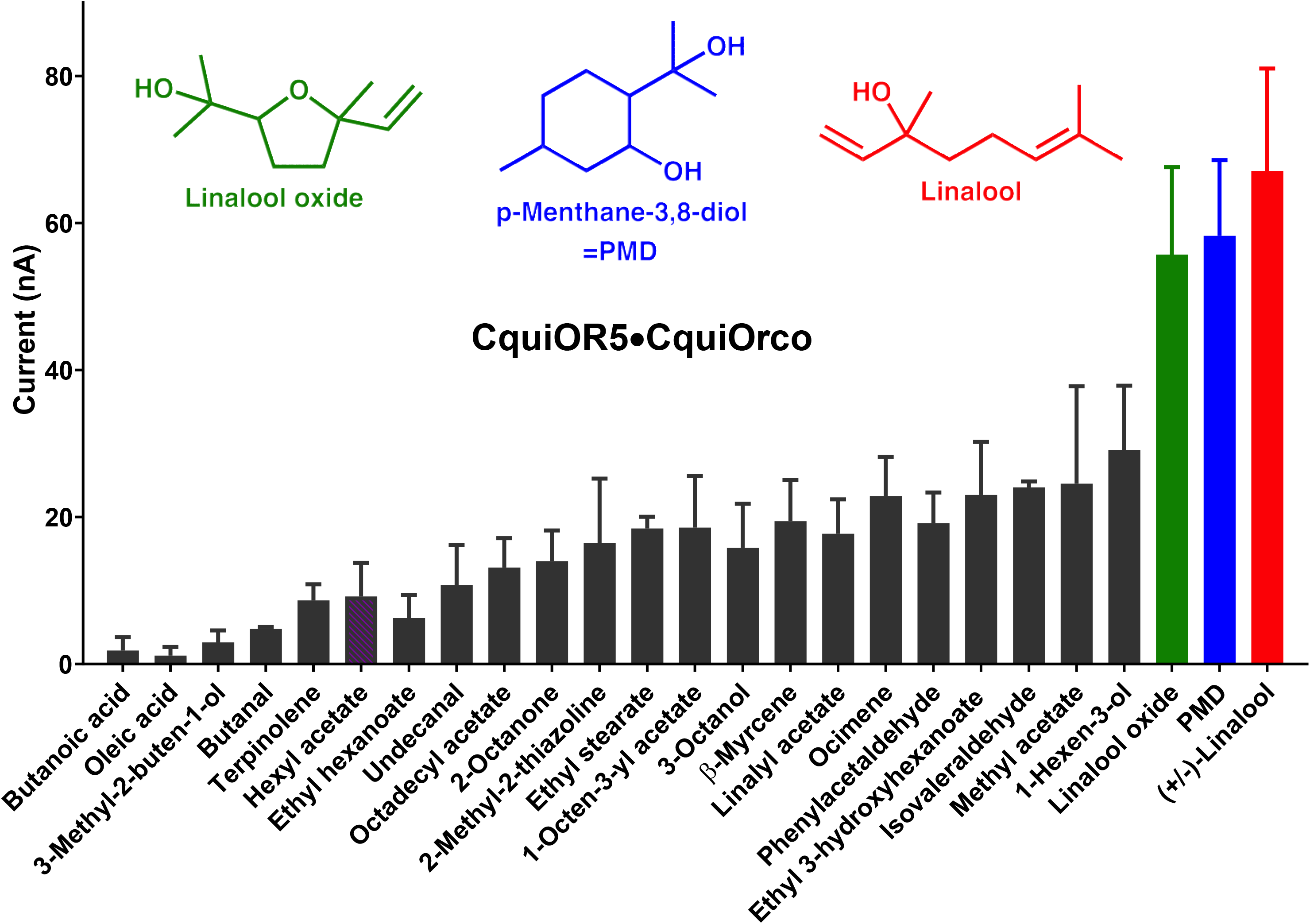
Quantification of current responses from oocytes expressing CquiOR5 and CquiOrco. All compounds in our panel were tested at 1 mM dose (n = 3). For clarity, bars are displayed in increasing order of response (Mean ± SEM) and by omitting compounds that did not elicit a response. The chemical structures of the three major ligands are displayed on the top of the graphic in colors corresponding to their bars.

The two receptors from *Ae. aegypti* in the cluster, ie, AaegOR14 and AaegOR15, gave remarkably different responses in terms of profile and sensitivity. AaegOR14/AaegOrco-expression oocytes elicited only weak currents when challenged with 4-methylphenol and 4-ethylphenol (Fig. S3C). By contrast, AaegOR15/AaegOrco-expressing oocytes generated dose-dependent, robust currents when challenged with phenethyl proprionate, phenethyl formate, and acetophenone (Fig. 4). As far as the two major ligands are concerned, AaegOR15 profile resembles that of CquiOR4 + CquiOrco.

**Fig. 4.**
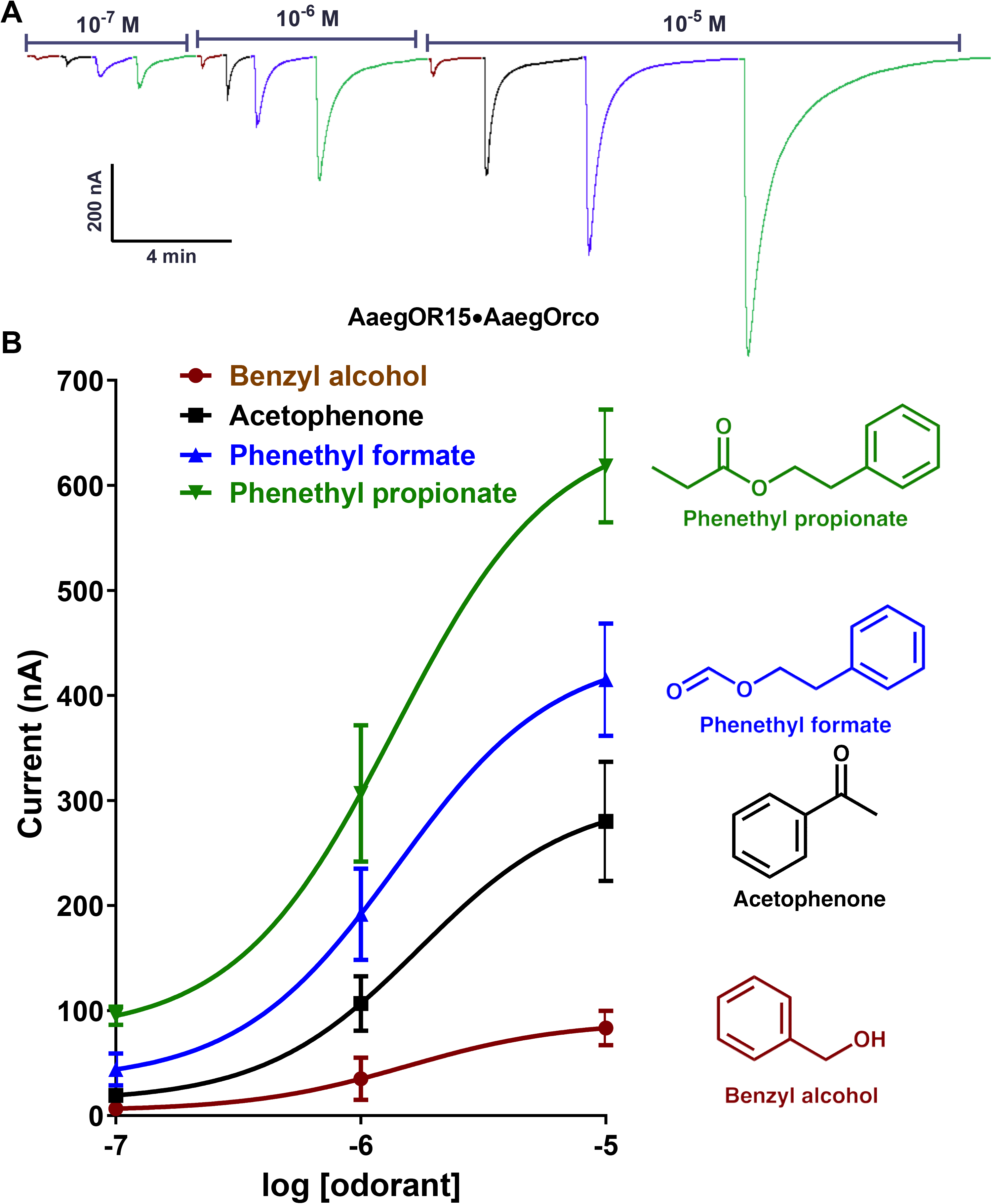
Concentration-dependent responses of AaegOR15⋅AaegOrco-expressing oocytes to four major odorants. (A) Traces obtained with a single oocyte challenged with phenethyl propionate (green), phenethyl formate (strawberry), acetophenone (black) and benzyl alcohol (cayenne) at the doses of 10^−7^ to 10^−5^ M. (B) Dose-dependent responses obtained with three oocytes with three replicates for each ligand. Responses (Mean ± SEM) were not normalized.

### 3.2. Tissue Expression Analysis

The best ligands for CquiOR4 and CquiOR5 have been previously reported to have repellency activity, particularly the commercially available PMD and 2-phenylethanol (USDA, 1947). To get a better insight into the possible role of these receptors in repellency behavior, we performed qPCR analyses. We surmised that transcript levels of the receptor for repellents and host attractants would decrease after a blood meal, whereas transcripts of receptors involved in the reception of oviposition attractants would increase. Transcript levels of CquiOR4 decreased significantly (P = 0.0122) after a blood meal (Fig. 5A). Additionally, this receptor was highly enriched in female antennae, with only basal levels in maxillary palps, proboscis and legs (Fig. 5A) thus further supporting a possible role in hosting finding or repellency activity. By contrast, transcript levels of CquiOR5 did not change significantly after a blood meal (Fig. 5B). Moreover, this receptor is not specific to or enriched in female antennae. We found high transcript levels in the proboscis, maxillary palps, and legs (Fig. 5B).

**Fig. 5.**
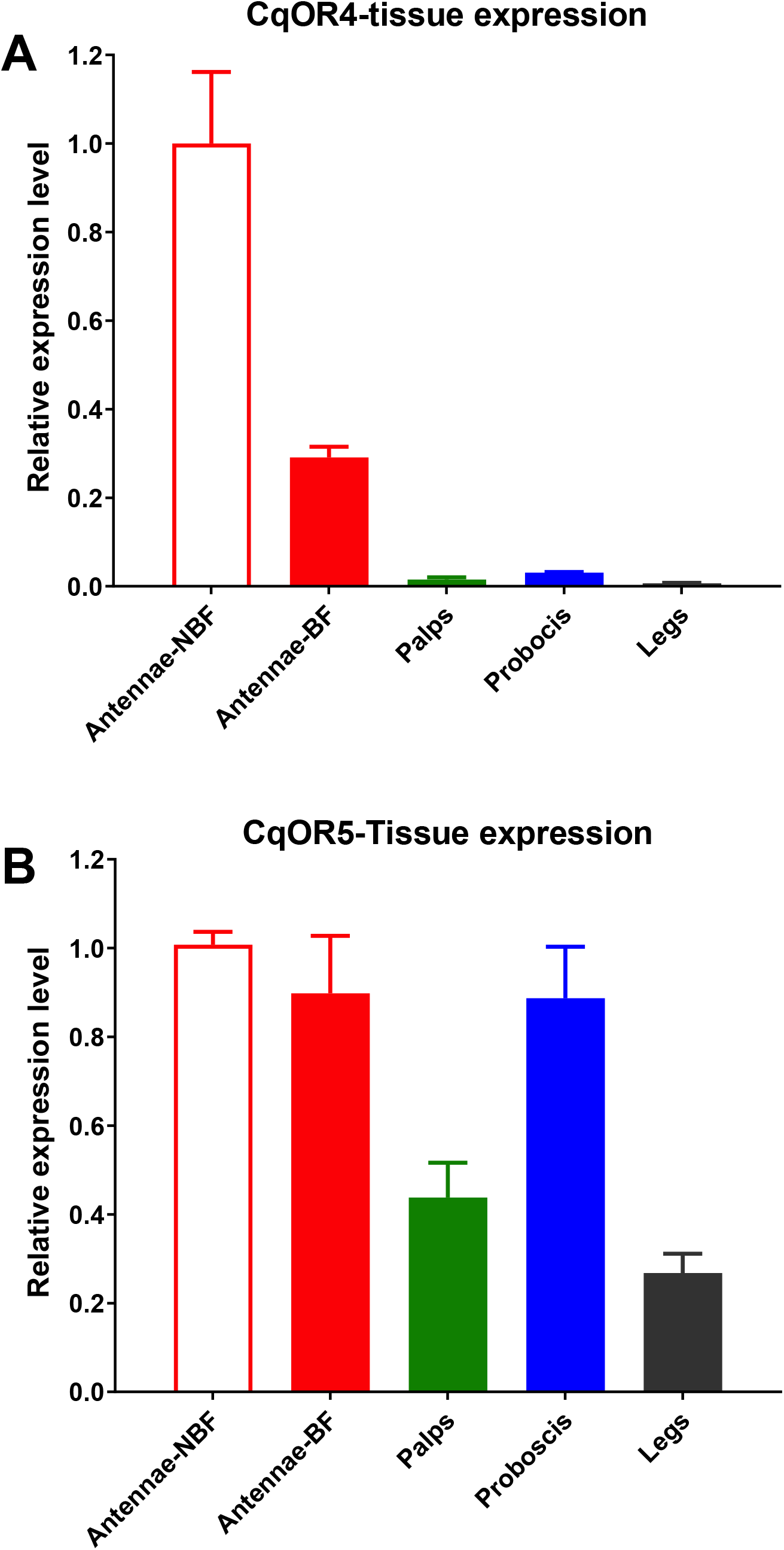
Transcript levels of *CquiOR4* and *CquiOR5* in *Cx. quinquefasciatus* tissues. (A) *CquiOR4* transcript levels in antennae of nonblood fed (NBF) and blood-fed (BF) female mosquitoes, maxillary palps, proboscis, and legs (specifically, tarsi from fore-, mid-, and hindlegs). (B) *CquiOR5* transcript levels in the same tissues described above (A). Data were obtained with three biological replicates, each with three technical replicates. Each biological replicate was normalized to the levels of expression in antennae of nonblood fed females.

### 3.3. Repellency Assays

Previously, 2-phenylethanol has been tested against the yellow fever mosquito (USDA, 1947) and reported to be active between 61 to 120 min. Using our surface landing and feeding assay (Leal et al., 2017; Xu et al., 2014), we tested repellency activity against the Southern house mosquito. In our preliminary assessment, 0.01% 2-phenylethanol gave 34.5±6.1 % protection (n =8), whereas at 0.1% it provided significantly higher protection 66.8±2.7 % (n = 7, Mann-Whitney test, P= 0.0003). We then compared 2-phenylethanol-elicited protection with that obtained with DEET, both at 1% and they showed comparable activity: DEET, 98.7±0.9%; 2-phenylethanol, 96.1±2.1% (n = 10 each; Mann-Whitney test, P = 0.6313). It is noteworthy mentioning that 2-phenylethanol is more volatile than DEET; thus, it may appear as strong as DEET, but may lose activity more rapidly, consistent with the earlier tested against *Ae. aegypti* (USDA, 1947). One of the main features of DEET is a long complete protection time.

### 3.4. Attempts to Link Odorant Reception with Repellency Activity

We performed RNAi experiments to test whether reducing transcript levels of *CquiOR4* would affect repellency activity. Similarly, we tested CquiOR5 vis-à-vis the best ligands. First, we compared the transcript levels of these genes in water-, β-galactosidase-dsRNA- and CquiOR4-dsRNA-injected mosquitoes. Transcript levels of *CquiOR4* decreased significantly (P = 0.0003, unpaired, two-tailed t-test) in knockdown mosquitoes (Fig. S4A). Similarly, CquiOR5-dsRNA-treated mosquitoes had significantly lower transcripts of *CquiOR5* than mosquitoes injected with water (P = 0.0013, unpaired, two-tailed t-test) or β-galactosidase-dsRNA.

Next, RNAi-treated mosquitoes were used to compared repellency activity. Because RNAi treatment reduced transcript levels only by ca. 60%, it is important to test repellency at low doses, otherwise, a possible link between reception and behavior may be overlooked. When tested at 0.01%, protection conferred by 2-phenylethanol, albeit low, was significantly reduced (n = 12-15, Mann-Whitney test, P=0.0062) (Fig. 6A). At a higher dose (0.1%), 2-phenylethanol-elicited protection is lower in CquiOR4-dsRNA-treated than in β-galactosidase-dsRNA-treated mosquitoes, but there was no significant difference (n = 6-7, Mann-Whitney test, P = 0.2191). By contrast, DEET-elicited repellency in these two groups of mosquitoes was not significantly different (n = 4, Mann-Whitney test, P > 0.999) (Fig. 6A).

**Fig. 6.**
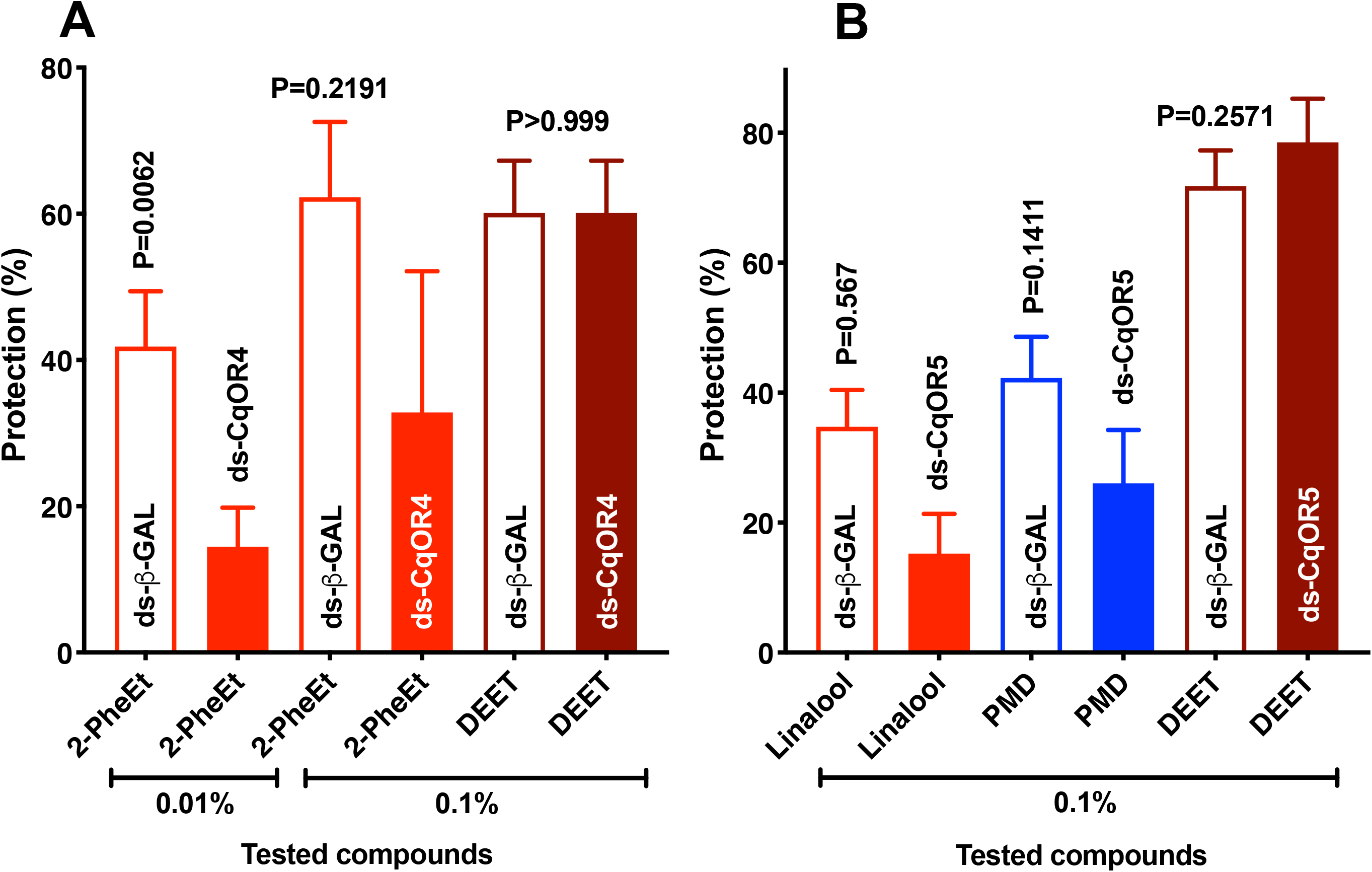
Graphical representation of repellency assays. For comparing response under different conditions, data were transformed to percent protection followed by unpaired comparisons. (A) Effect of CquiOR4 on the response of *Cx. quinquefasciatus* to 2-phenhylethanol and DEET. At a very low dose (0.01%), 2-phenylethanol showed a low protection rate, which was significantly reduced in CquiOR4-dsRNA- as compared to β-galactosidase-dsRNA-treated mosquitoes. At a higher dose (0.1%), 2-phenylethanol-elicited protection was higher, but also significantly reduced in RNAi-treated mosquitoes. By contrast, RNAi treatment did not affect DEET-elicited repellency (positive control). n= 6-15; for DEET n = 4. (B) Effect of CquiOR5 on *Cx. quinquefasciatus* responses to linalool, PMD, and DEET at 0.1%. When comparing repellency activities elicited by linalool and PMD in CquiOR5-dsRNA- with β-galactosidase-ds-RNA-injected mosquitoes, a nonsignificant reduction was observed, thus suggesting that other ORs might be involved. n = 11-16; for DEET n = 11. All statistical comparisons were performed with two-tailed Mann-Whitney Test.

Measuring linalool-elicited repellency activity at 0.1% showed a non-significant reduction of protection in CquiOR5-dsRNA-treated mosquitoes as compared to β-galactosidase-dsRNA-treated mosquitoes (n = 11-16, Mann-Whitney test, P = 0.567) (Fig. 6B). Likewise, protection by PMD was also reduced, but not significantly different (n=12-16, Mann-Whitney test, P = 0.1411) (Fig. 6B). There was no significant difference in repellency (protection) elicited by DEET (n = 12, Mann-Whitney test, P = 0.2571) (Fig. 6B). Attempts to test repellency at lower doses were unrewarding. At 0.05% linalool, 56±5.1% of mosquitoes responded to the control side of the arena, whereas 44±5% responded to treatment (n =6). Likewise, PMD at 5% provided no protection: control, 51.7±2%; treatment, 48.2±2%. Thus, we were unable to test the effect of RNAi treatment on repellency activity at lower than 0.1% dose.

In conclusion, repellency activity mediated by linalool and PMD might involve multiple receptors. Here, we show that PMD activates not only CquiOR5, but also CquiOR84/85. Previously, we showed that PMD activated a DEET receptor in the Southern house mosquito, CquiOR136 (Xu et al., 2014). The reduction in protection observed with knockdown mosquitoes, albeit not statistically significant, suggests that CquiOR5 might be one of the receptors mediating linalool and PMD repellency activity. On the other hand, we cannot rule out the possibility that other receptors are involved in a combinatorial code reception of 2-phenylethanol, but the significant reduction in protection in CquiOR4-dsRNA-treated mosquitoes suggests it may play a significant part in 2-phenylethanol-mediated repellency activity.

## Acknowledgments

The authors are grateful to Dr. Anthon J. Cornel, UC Davis, Department of Entomology and Nematology, for providing *Ae. aegypti* mosquitoes for RNA extraction. We thank all laboratory members for discussions during our weekly seminars and Dr. Su Liu for comments on an earlier draft of the manuscript.

## Author Contributions

F. Z., P.X. and W.S.L. designed the experiments and performed data analysis. W.S.L. conceived the project. F. Z. and P.X. carried out molecular biology and electrophysiology work. F.Z. performed RNAi experiments and measured mosquito behavior. W.S.L. prepared figures and wrote the manuscript. All authors provided input, read, and approved the final version of the manuscript.

---

**Fig. S1.**
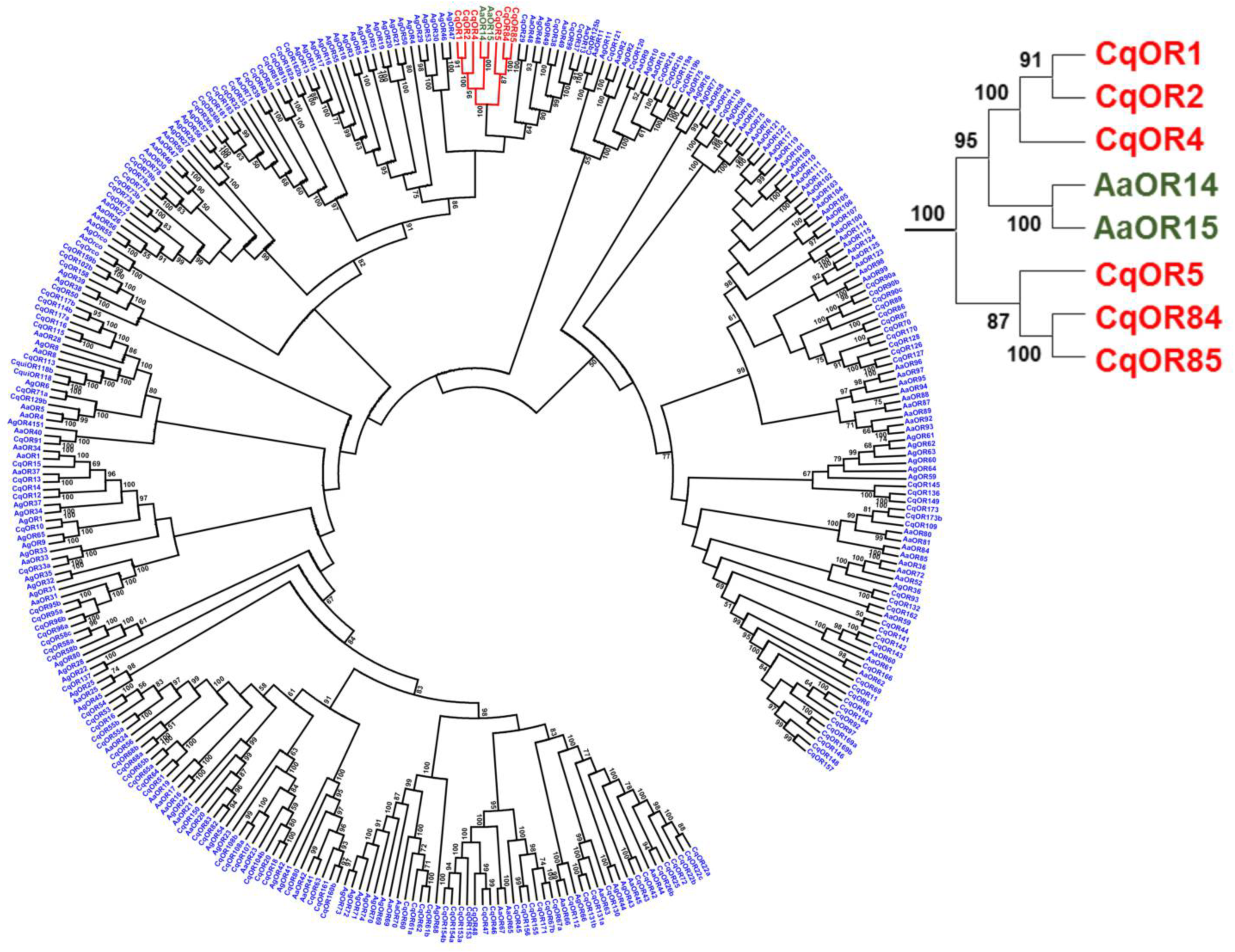
Phylogenetic relationships of ORs from *Cx. quinquefasciatus* (Cq), *Aedes aegypti* (Aa), and *Anopheles gambiae* (Ag). The evolutionary history was inferred using the Neighbor-Joining method (Saitou and Nei, 1987). The percentage of replicate trees in which the associated taxa clustered together in the bootstrap test (1000 replicates) are shown next to the branches (Felsenstein, 1985). The tree is drawn to scale, with branch lengths in the same units as those of the evolutionary distances used to infer the phylogenetic tree. The evolutionary distances were computed using the p-distance method (Nei and Kumar, 2000) and are in the units of the number of amino acid differences per site. The analysis involved 330 amino acid sequences. All positions containing gaps and missing data were eliminated. There was a total of 264 positions in the final dataset. Evolutionary analyses were conducted in MEGA7 (Kumar et al., 2016). The tree is available in iTOL: https://itol.embl.de/tree/169237242541921536268595

**Fig. S2.**
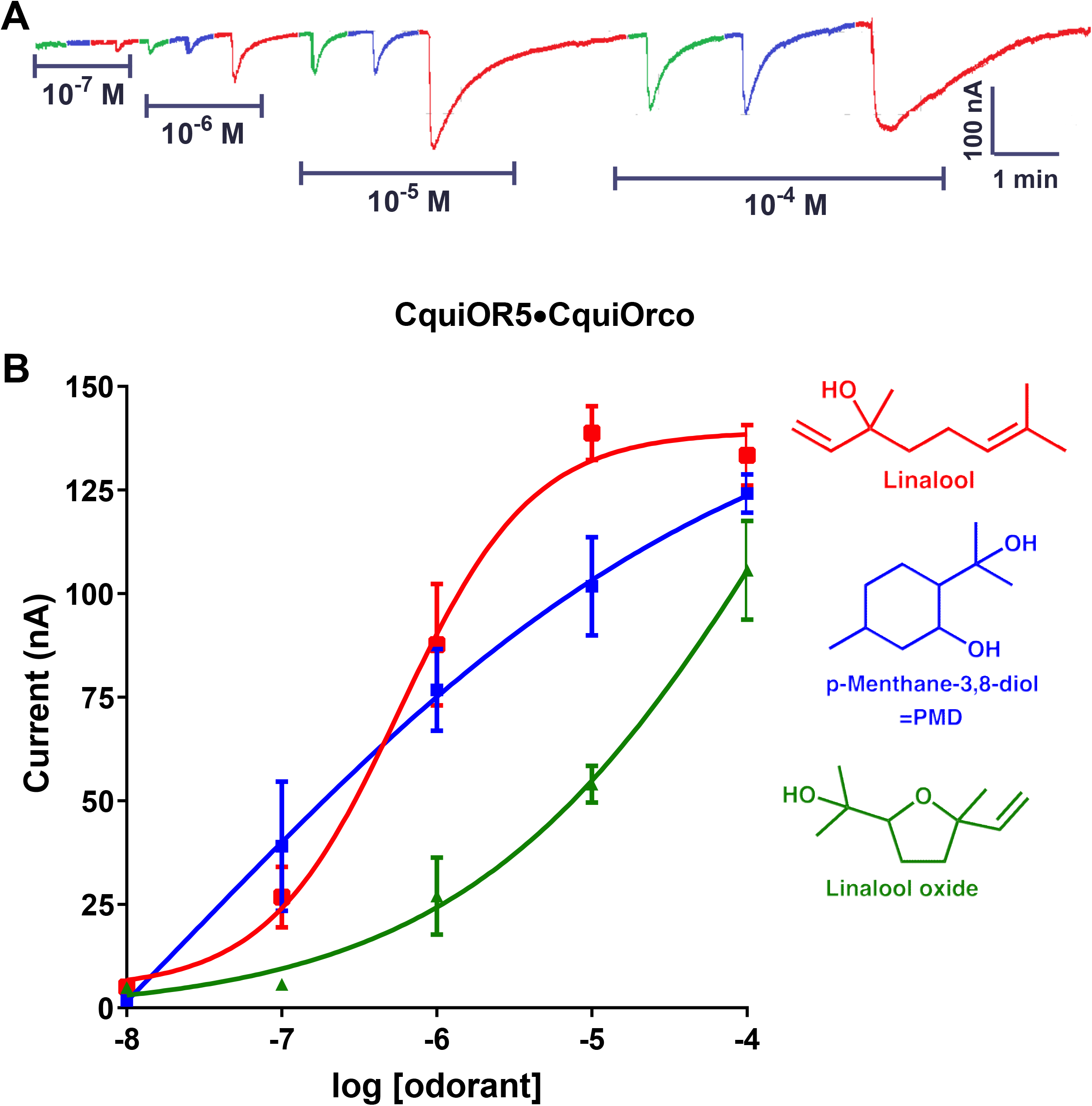
Concentration-dependent responses of CquiOR5⋅CquiOrco-expressing oocytes to three major odorants. (A) Traces obtained with a single oocyte challenged with racemic linalool (red), PMD (strawberry) and linalool oxide (green) at the doses of 0.1 µM (10^−7^ M) to 0.1 mM (10^−4^ M). (B) Dose-dependent responses obtained with three oocytes and three replicates for each ligand. Responses (Mean ± SEM) were not normalized.

**Fig. S3.**
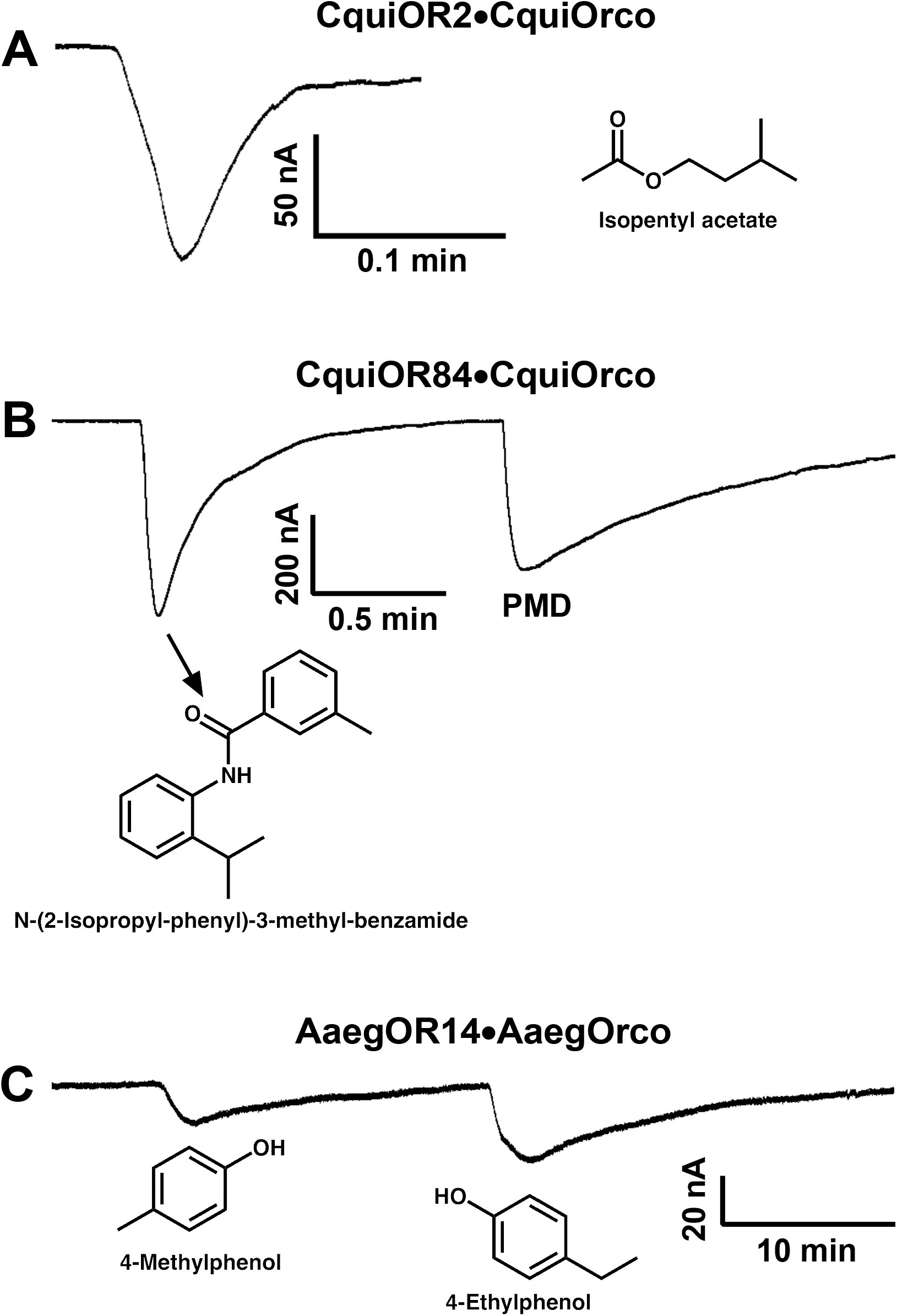
Currents elicited by compounds in our test panel on oocytes expressing the specified ORs. (A) Response of CquiOR2⋅CquiOrco-expressing oocytes to isopentyl acetate. (B) Current elicited by N-(2-isopropyl-phenyl)-3-methyl-benzamide and PMD on oocytes expressing CquiOR84 and CquiOrco. (C) Responses elicited by 4-methylphenol and 4-ethylphenol on AaegOR14⋅AaegOrco-expressing oocytes. These responses were repeated at least with two different oocytes. Note the weak responses in A and C.

**Fig. S4.**
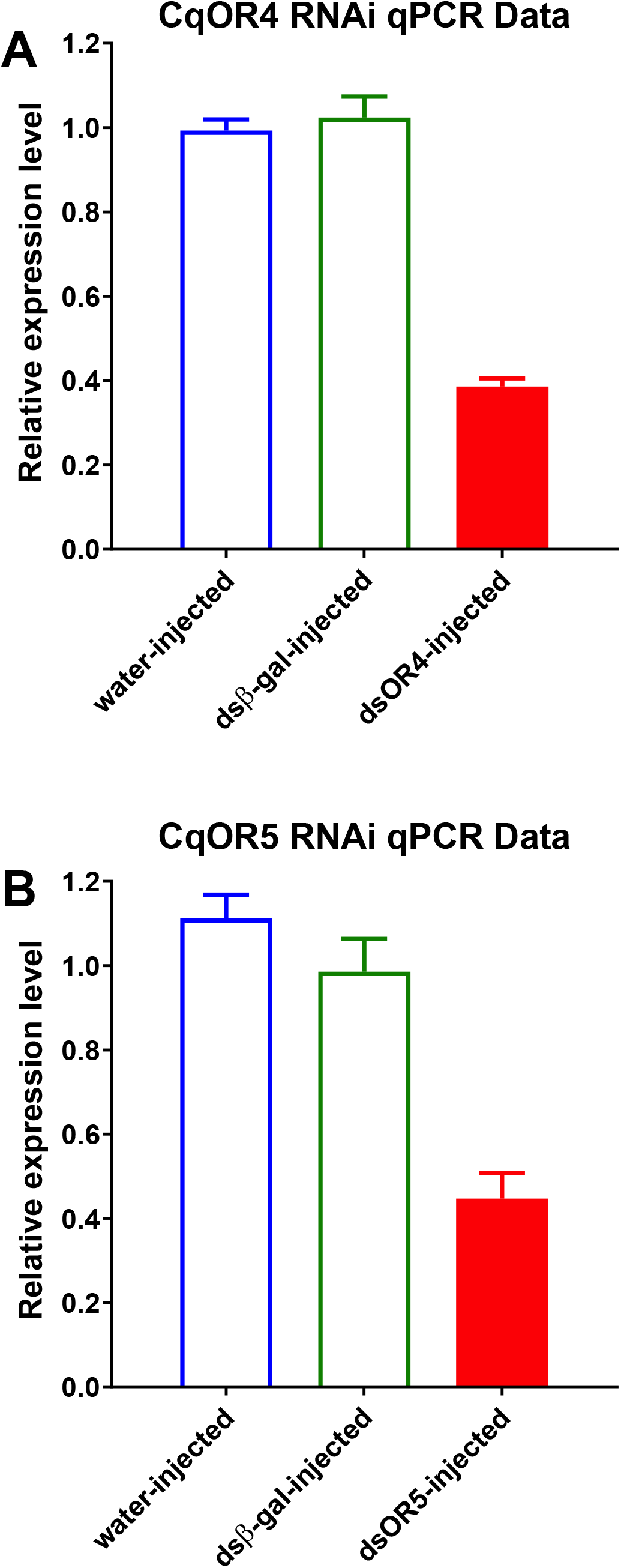
qPCR data for evaluation of RNAi treatment. (A) Transcript levels of CquiOR4 were significantly reduced (P=0.0003) compared to both water-injected mosquitoes and those treated with β-galactosidase-dsRNA. (B) Mosquitoes injected with CquiOR5-dsRNA had significantly lower (P=0.0013) transcript levels of CquiOR5 when compared to both mosquitoes treated with dsRNA of a control gene and those injected with water.

